# AATF-mediated Liver Damage and Inflammation to Cancer: Therapeutic Intervention by Curcumin in Experimental MASH-HCC

**DOI:** 10.1101/2023.05.26.542402

**Authors:** Akshatha N. Srinivas, Diwakar Suresh, Saravana Babu Chidambaram, Prasanna K. Santhekadur, Divya P. Kumar

**Affiliations:** Department of Biochemistry, CEMR lab, JSS Medical College, JSS Academy of Higher Education and Research, Mysuru, Karnataka, India; Department of Pharmacology, JSS College of Pharmacy, JSS Academy of Higher Education and Research, Mysuru, Karnataka, India

**Keywords:** Curcumin, metabolic dysfunction associated steatohepatitis, hepatocellular carcinoma, apoptosis antagonizing transcription factor, inflammation, tumorigenesis

## Abstract

In tandem with the expanding obesity pandemic, the prevalence of metabolic dysfunction associated steatohepatitis (MASH, formerly known as NASH)-driven hepatocellular carcinoma (HCC) is predicted to rise globally, creating a significant need for therapeutic interventions. We previously identified the upregulation of apoptosis antagonizing transcription factor (AATF), which is implicated in facilitating the progression from MASH to HCC. The objective of this study was to examine whether the intervention of curcumin could alleviate AATF-mediated MASH, inhibit tumor growth, and elucidate the underlying mechanism. A preclinical murine model mimicking human MASH-HCC was employed, subjecting mice to either a chow diet normal water (CDNW) or western diet sugar water (WDSW) along with very low dose of carbon tetrachloride (CCl_4_-0.2 μl/g, weekly). Mice receiving curcumin (CUR) alongside WDSW/CCl_4_ exhibited significant improvements, including reduced liver enzymes, dyslipidemia, steatosis, inflammation, and hepatocellular ballooning. Curcumin treatment also suppressed hepatic expression of inflammatory, fibrogenic, and oncogenic markers. Of note, there was a significant reduction in the expression of AATF upon curcumin treatment in WDSW/CCl_4_ mice and human HCC cells. In contrast, curcumin upregulated Kruppel-like factor 4 (KLF4) in MASH liver and HCC cells, which is known to downregulate sp1 (specificity protein-1) expression. Thus, curcumin treatment effectively inhibited the progression of MASH to HCC by downregulating the expression of AATF via the KLF4-Sp1 signaling pathway. These preclinical findings establish a novel molecular connection between curcumin and AATF in reducing hepatocarcinogenesis, and provide a strong rationale for the development of curcumin as a viable treatment for MASH-HCC in humans.

## Introduction

Hepatocellular carcinoma (HCC), a highly aggressive type of liver cancer known to develop in the setting of chronic liver disease, is a serious health care challenge worldwide [1]. There is mounting evidence that the etiologies of HCC are rapidly shifting towards obesity and the metabolic syndrome [2]. Given the 25% global prevalence of metabolic dysfunction associated steatotic liver disease (MASLD, formerly known as NAFLD), a hepatic manifestation of metabolic syndrome, MASLD-related HCC is an emerging cause of liver tumorigenesis [3,4]. The metabolic dysfunction associated steatohepatitis (MASH, formerly known as NASH), a progressive variant of MASLD, increases the likelihood of developing cirrhosis and HCC. The exact mechanisms by which MASH leads to HCC are not fully understood, but it is believed that chronic inflammation and cellular damage caused by MASH accelerate fibrogenesis and create an environment that is more conducive to cancerous cell growth and development [5]. MASH-associated HCC is often associated with poor survival due to several factors, including poor prognosis, late diagnosis, and inadequate management. Unlike other types of HCC, MASH-HCC can develop even in the absence of cirrhosis, making it more difficult to detect and manage [6]. The complexities and heterogeneity of the MASH-HCC pathophysiology have hampered treatment strategies. Multifactorial processes, including genetic, metabolic, immunogenic, and endocrine mechanisms, contribute to the transition of damaged hepatocytes into HCC [7,8,9]. At a cellular level, genetic and epigenetic events orchestrate the pro-oncogenic milieu against a background of insulin resistance and inflammation [10,11]. Thus, a mechanistic approach to studying the root cause of MASH-HCC and the development of therapeutic strategies for MASH-HCC is the need of the hour.

Apoptosis antagonizing transcription factor (AATF) is a multifunctional protein known to play a critical role in key cellular processes [12]. AATF is an emerging modulator of cell proliferation, survival, and apoptosis depending on the cellular status, thereby serving as both a prooncogene and a tumor suppressor gene [13–15]. Studies have shown the involvement of the AATF in many cancers, such as leukemia, breast cancer, neuroblastoma, lung cancer, and colon cancer [16–20]. We have previously shown elevated levels of hepatic AATF in human and mouse MASH-HCC [21]. In a diet-induced animal model of NAFLD (DIAMOND), AATF contributes to tumorigenesis via STAT3-dependent activation of monocyte chemoattractant protein-1 (MCP-1) [21]. AATF is also known to significantly contribute to various hallmarks of HCC, and we have recently elucidated the regulatory role of AATF in HCC angiogenesis [22].

Natural monomeric products are extensively studied for their hepatoprotective and anti-inflammatory efficacy, which can improve MASH [23]. Curcumin, an active component of curcuminoids derived from the rhizome of turmeric (Curcuma longa, Zingiberaceae), is a diarylheptane and is known for its anti-inflammatory, antioxidant, and anti-cancer properties [24–26]. Curcumin is known to mediate anticancer effects by modulating inflammatory cytokines, growth factors, transcription factors, and multiple signaling proteins [27]. Several studies have shown the positive effects of curcumin on ameliorating MASH. Curcumin lowers de novo lipogenesis by reducing the levels of sterol regulatory element binding protein 1-c (SREBP-1c) and adipose differentiation-related protein (ADRP) [28]. Further, tumor necrosis factor-α (TNF-α) and reactive oxygen species (ROS) levels were also found to be reduced upon curcumin treatment in HFD-induced MASH [29]. In the streptozotocin (STZ)-induced MASH-HCC model, curcumin protected liver damage from ER stress and related inflammation by inhibiting the translocation of high-mobility group box 1 (HMGB1) and nuclear factor kappa B (NF-κB) [30]. However, the investigation on the specific effect of curcumin on AATF-mediated MASH-HCC and its anti-cancer mechanism in the context of inflammatory milieu remains unexplored.

The overall goal of the present study was to investigate if curcumin intervention could alleviate AATF-mediated MASH-associated liver tumorigenesis and elucidate its underlying mechanism. To achieve this, we employed the experimental MASH-HCC model, which is a fully characterized murine model that recapitulates the disease progression from steatohepatitis to fibrosis and HCC and closely resembles the metabolic, histological, and transcriptomic signature of human MASH-HCC. Using this model, we demonstrated that curcumin treatment attenuates the progression of MASH to HCC by ameliorating inflammation and fibrosis. In terms of the underlying mechanisms, our data suggest that curcumin exerts preventive effects on HCC development by suppressing AATF expression via kruppel-like factor 4 (KLF4) and the specificity protein 1 (Sp1) signaling pathway. Collectively, these findings offer novel insights into the hepatoprotective action of curcumin and its therapeutic significance in addressing MASH-associated HCC, mediated by AATF.

## Materials and Methods

### Materials

Curcumin, carboxy methyl cellulose (CMC), TRIzol, RIPA buffer, Sirius Red stain, corn oil, and AATF antibody were procured from Sigma-Aldrich (St. Louis, USA). Dulbecco’s modified eagle’s medium (DMEM), hematoxylin, DMSO, glucose, and fructose were procured from HiMedia Laboratories (India). Lipofectamine 3000, penicillin/streptomycin, glutamine, and FBS were from Invitrogen (USA); antibodies to Sp1, pJNK, pErk1/2, JNK, Erk1/2, and control and AATF shRNA from Santa Cruz Biotechnology, Inc. (CA, USA); antibodies to KLF4, Ki67, pSTAT3, STAT3, and β-actin from Cell Signaling Technologies (USA); verso cDNA synthesis kit, and DyNamo Colorflash SYBR green kit from Thermo Fisher Scientific (USA); western bright ECL HRP substrate from Advansta (USA); western blotting materials were from Bio-Rad; primers from Integrated DNA Technologies (IDT) (Iowa, USA), water-soluble tetrazolium salt (WST-1) (Takara, India). The western diet was customized and purchased from Research Diets Inc. (NJ, USA). The ALT, AST, and Total cholesterol kits were from Monlab tests (Spain) and all other reagents were obtained from Thermo Fisher Scientific or Sigma.

### Animals

Male C57Bl/6NCrl mice (6-7 weeks old) were purchased from Hylasco Biotechnology Pvt. Ltd. (Charles River Laboratories) and maintained as described previously [31]. All the mice were housed in a 12-hour light and 12-hour dark cycle in the animal facility of the Center for Experimental Pharmacology and Toxicology, JSS AHER. All the procedures were performed according to the protocol approved by the Committee for the Purpose of Control and Supervision of Experiments on Animals (CPCSEA), Govt. of India (JSSAHER/CPT/IAEC/060/2021).

### Study design and interventions

Following the one-week acclimatization, mice were fed either a chow diet (CD, lab diet, USA) with normal water (NW) or a high fat, high sucrose, and high cholesterol (WD, 21% fat, 41% sucrose, and 1.25% cholesterol by weight, Research Diet, Inc., USA) diet with sugar water (SW, 18.9 g/L d-glucose and 23.1 g/L d-fructose) ad libitum for the first 12 weeks. Simultaneously with dietary intervention, all the mice were treated with CCl_4_ at a dose of 0.2 μl/g of body weight dissolved in corn oil and administered intraperitoneally once a week throughout the study as described previously [32]. After 12 weeks, mice were randomly divided into four groups: CDNW with vehicle (0.5% CMC), CDNW with curcumin (100 mg/kg/day), WDSW with vehicle (0.5% CMC), and WDSW with curcumin (100 mg/kg/day) (**Figure 1A**). Curcumin and vehicle were administered every day for 12 weeks via oral gavage. Mice were continuously monitored for food intake, water intake, and body weight. After 24 weeks, mice were euthanized, and blood and tissues were collected for further analysis.

**Figure 1.**
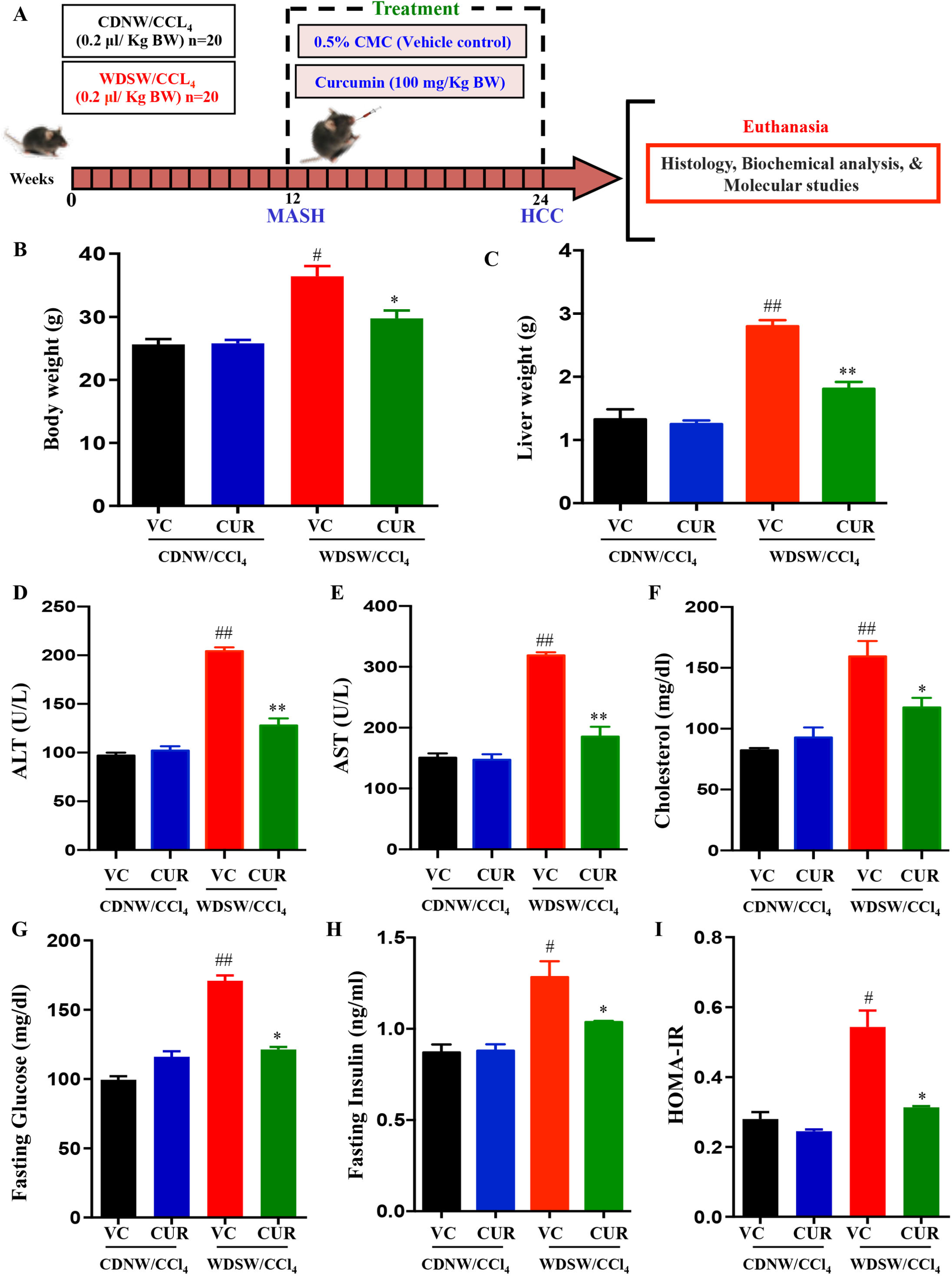
Curcumin treatment reduced obesity and improved liver injury and insulin resistance in WDSW/CCl_4_ mice. Mice were given either CDNW or WDSW for 12 weeks before being divided into two further groups and treated with either vehicle (VC) or curcumin (CUR) for the next 12 weeks (A). Before euthanasia, mice were measured for (B) body weight and (C) liver weight. Blood collected from overnight fasted mice was used for determining (D) ALT, (E) AST, and (F) total cholesterol. (G) Fasting glucose and (H) fasting insulin were measured, and (I) HOMA-IR was calculated using the formula (fasting insulin (milliunits/liter) - fasting glucose (mmol/liter))/22.5. All the data are expressed as mean±SEM with 6-8 mice in each group. ## p < 0.001 or # p < 0.05 compared to CDNW/CCl_4_ with vehicle control; ** p < 0.001 or * p < 0.05 compared to WDSW/CCl_4_ with vehicle control.

### Biochemical analysis

The blood collected via the retro-orbital puncture route from the overnight fasted mice was processed by centrifuging at 1500 g for 15 minutes at 4°C. The serum collected was stored at −80°C. Serum ALT, AST, and Total cholesterol were measured according to the manufacturer’s instructions [33].

### Liver Histology

Formalin-fixed, paraffin-embedded liver tissues were stained with hematoxylin and eosin (H&E) staining to evaluate liver histology for steatosis, lobular inflammation, and hepatocyte ballooning. MASLD activity scoring was evaluated by an expert pathologist (double blindfolded) at JSS Hospital according to the FLIP algorithm and NASH-Clinical Research Network (CRN) criteria [34,35].

### Sirius Red Staining

The formalin-fixed, paraffin-embedded liver tissue sections were deparaffinized and rehydrated using xylene and a series of ethanol concentrations. Slides were stained with hematoxylin for 10 minutes. Following a water wash, the slides were stained by Sirius red solution for 1 hour at room temperature. The excess stain was removed by washing it in 0.5% acidified water for 5 minutes. Slides were mounted with coverslips, and the images were taken with an Olympus BX53 microscope and quantified using the ImageJ software.

### Immunostaining

The formalin-fixed, paraffin-embedded liver tissue sections were deparaffinized and rehydrated using xylene and a series of ethanol concentrations. Slides were incubated with citrate buffer (pH 6) for 15 minutes at 94°C for antigen retrieval, followed by a water wash and incubation with 3% hydrogen peroxide for 10 min. After the incubation with blocking buffer for 1 h at room temperature, sections were added with the target of interest and incubated overnight at 4°C in a humidified chamber. Polyexcel HRP/DAB detection system-one step (PathnSitu, Biotechnologies) was used for developing the signals and hematoxylin for counterstaining the nucleus. Antibody dilution for AATF (1:100) and Ki67 (1:200). All the images were taken using an Olympus BX53 microscope and quantified using the ImageJ software.

### RNA isolation and quantitative real-time PCR

Total RNA was isolated from the cells and liver tissues using the TRIzol reagent according to the manufacturer’s protocol. The RNA was quantified using a nanodrop spectrophotometer and reverse transcribed using a Verso cDNA synthesis kit. Gene expression analysis was carried out in the Rotor-Gene Q 5plex HRM System (QIAGEN) using the DyNamo Colorflash SYBR Green kit, 0.5 mM primers, and 50 ng of cDNA in a 20 μl reaction volume. The relative fold change in gene expression was calculated as 2^-ΔΔCt^ and relative fold changes were calculated by normalizing the values to internal control, β actin. The validated primer sequences used are as follows: mouse IL-1β (Forward primer 5′- TGCCACCTTTTGACAGTGATGA-3′ and Reverse primer 5′TGATGTGCTGCTGCGAGATTTG-3′); mouse IL-6 (Forward primer 5′- GGAGCCCACCAAGAACGATA-3′ and Reverse primer 5′- TGCCACCTTTTGACAGTGATGA-3′ and Reverse primer 5′TGATGTGCTGCTGCGAGATTTG-3′); mouse IL-6 (Forward primer 5′- GGAGCCCACCAAGAACGATA-3′ and Reverse primer 5′- AACTGGATGGAAGTCTCTTGC-3′); mouse TNF-α (Forward primer 5′- TAGCCCACGTCGTAGCAAACC-3′ and Reverse primer 5′- CTTTGAGATCCATGCCGTTGGC-3′); mouse α-SMA (Forward primer 5′- CTACTGCCGAGCGTGAGATTGT-3′ and Reverse primer 5′- CCCGCTGACTCCATCCCAATGA-3′); mouse TGF-β (Forward primer 5′- GCTGCATATCGTCCTGTGG-3′ and Reverse primer 5′- CTTCCATTTCCACATCCGACT-3′); mouse CD31 (Forward primer 5′- GTGGAAGTGTCCTCCCTTGA-3′ and Reverse primer 5′- GGGAGCCTTCCGTTCTAGAGTAT-3′); mouse AFP (Forward primer 5′- AACCTCCAGGCAACAACCATT-3′ and Reverse primer 5′- CACTCCTCGTGGATGTGAGC-3′); human AATF (Forward primer 5′- TAGAACGGAAGACCAGCTCC-3′ and Reverse primer 5′- TGCTAAGGACATGAAACCGA-3′, mouse AATF (Forward primer 5′- GAGTGATGATGCCAGGACGGA-3′ and Reverse primer 5′- ACTGTCACTTCCCACGGTCTG-3′); human KLF4 (Forward primer 5′- GATGCTCACCCCACCTTCTTC-3′ and Reverse primer 5′- GTAAGGTTTCTCACCTGTGTGG-3′, mouse KLF4 (Forward primer 5′- CACCTGGCGAGTCTGACAT-3′ and Reverse primer 5′- TTCCTCACGCCAACGGTTA-3′); human Sp1 (Forward primer 5′- CCACCATGAGCGACCAAGAT-3′ and Reverse primer 5′- AAGGCACCACCACCATTACC-3′; mouse Sp1 (Forward primer 5′- GCCACCATGAGCGACCAAGAT-3′ and Reverse primer 5′- GCCAGCAGAGCCAAAGGAGAT-3′); human β-actin (Forward primer 5′- AGAGATGGCCACGGCTGCTT -3′ and Reverse primer 5′- CAGGACTCCATGCCCAGGAA-3′; mouse β-actin (Forward primer 5′- CAGCCTTCCTTCTTGGGTATGG-3′ and Reverse primer 5′- CCTGCTTGCTGATCCACATCT-3′)

### Immunoblotting

Fresh lysates were prepared from the cells and liver tissues using RIPA buffer containing protease/phosphatase inhibitors. The supernatants were collected after centrifuging the homogenate at 13000 rpm for 10 minutes at 4°C. The protein concentration of the lysates was determined by Bradford’s protein estimation method. An equal amount of lysates (30 μg) were loaded into SDS-PAGE to separate the protein and transferred onto a nitrocellulose membrane. Further, the membranes were blocked with 5% non-fat skim milk for 1 hour and incubated with the target primary antibodies (AATF, pJNK, JNK, pErk1/2, Erk1/2, KLF4, Sp1, pSTAT3, STAT3 and β-actin) overnight at 4°C. Following this, the membrane was incubated with a secondary antibody for 2 h at room temperature. Western Bright ECL HRP substrate was used to visualize the blots in the UVitec Alliance Q9 chemiluminescence imaging system. Images were analyzed using ImageJ software and bands were normalized with corresponding internal controls.

### Cell culture and generation of stable cell lines

Human HCC cell line - QGY-7703 (a kind donation from Dr. Devanand Sarkar, Virginia Commonwealth University, Richmond, USA) was cultured in Dulbecco’s modified eagle’s medium containing 4.5 g/L glucose and supplemented with 10% fetal bovine serum, L-glutamine, and 100 U/ml penicillin-streptomycin incubated at 37°C in 5% CO_2_. The authenticity of QGY-7703 cells was confirmed by short tandem repeat (STR) profiling.

In QGY-7703 cells, AATF shRNA expressing stable clones were generated as previously described [21]. Different concentrations of puromycin (1-10 μg/ml) were used to determine the optimal antibiotic concentration for selecting stable cell colonies. According to the manufacturer’s instructions, the control and AATF shRNA plasmids containing the puromycin resistance gene were transfected into QGY-7703 cells. Single colonies were then chosen and grown in 1 μg/ml puromycin. AATF knockdown was confirmed by both qRT-PCR and immunoblotting analysis.

### Cell viability assay

The effect of curcumin on cell viability was determined by the WST-1 assay. Briefly, 1×10^4^ cells were seeded into a 96-well plate and allowed to attain 70% confluency. Different concentrations of curcumin (0, 10, 20, 40, 60, 80, and 100 μM) dissolved in DMSO were treated in cells for 24 h, 48 h, and 72 h. The final concentration of DMSO in the treatment medium was < 0.05%. To assess the cell viability, control and AATF knockdown QGY 7703 cells were treated with curcumin (20 μM/ml) for 24 hours. At the end of the treatment, a 1:10 diluted WST-1 mix solution was added to each well and incubated for 60 minutes at 37°C in 5% CO_2_. The absorbance was measured at 450 nm on a multi-mode plate reader (EnSpire™ Multimode Plate Reader, Perkin Elmer). The data is represented as percent cell viability after normalizing with a negative control (no treatment).

### Migration assay

QGY 7703 control and AATF knockdown cells were seeded at a density of 5×10^5^ in a 6-well plate and cultured up to 70% confluency. A gentle scratch was done using a 1000 μl pipette tip across the center of each well. The detached cells were removed with PBS. Cells were incubated with either vehicle or curcumin (20 μM/ml) containing fresh media. After 24 h, cells were observed under the microscope for migration ability. The images were taken using a Zeiss Primovert inverted microscope, and the gap distance was measured using ImageJ software.

### Statistical analysis

Results were calculated as mean +/- SEM. Statistical significance was analyzed using Student’s t-test for two groups or analysis of variance (ANOVA) with post hoc Bonferroni correction for multiple comparisons. All statistical analyses were performed using the GraphPad Prism software (version 6), and p values < 0.05 (*/^#^) or < 0.001 (**/^##^) were considered significant.

## Results

### Curcumin attenuates the progression of MASH to HCC

To assess the effect of curcumin treatment on MASH-associated HCC, a 24-week study was performed on C57Bl/6 mice. Mice were divided into two groups and fed with either chow diet normal water (CDNW) or western diet sugar water (WDSW). Both groups received a low dose of CCl4 (0.2 μl/g body weight) intraperitoneally once a week until the end of the study. In the 12^th^ week, upon confirmation of MASH in the WDSW/CCl_4_ group, the mice were further divided into 4 groups: CDNW/CCl_4_ + vehicle control (VC), CDNW/CCl_4_ + curcumin (100 mg/kg body weight), WDSW/CCl_4_ + vehicle, and WDWW/CCl_4_ + curcumin (100 mg/kg body weight) (**Figure 1A**). The mice treated with vehicle in WDSW/CCl_4_ gained significant body weight and liver weight compared to CDNW/CCl_4_ mice (**Figure 1B and 1C**). This was consistent with hypercholesteremia and elevated liver enzymes in mice treated with vehicle in WDSW/CCl_4_, thereby confirming severe liver injury (**Figure 1D, 1E, and 1F**). Curcumin treatment in the WDSW/CCl_4_ mouse groups for 12 weeks showed a notable reduction in both body weight and liver weight compared to the vehicle treated WDWW/CCl_4_ mice (**Figure 1B and 1C**). Similarly, ALT, AST, and total cholesterol levels were also lowered by curcumin treatment in WDSW/CCl_4_ mice (**Figure 1D, 1E, and 1F**) when compared to CDNW/CCl_4_ mice. No effects were seen in CDNW/CCl_4_ mouse groups treated with curcumin. Additionally, the administration of curcumin also had considerable effects on fasting glucose and fasting insulin levels (**Figure 1G and 1H**), along with HOMA-IR in WDSW/CCl_4_ mice (**Figure 1I**). Thus, these data demonstrate that curcumin treatment ameliorated obesity, insulin resistance, and liver injury in the WDWW/CCl_4_ mouse groups.

Histological examinations were carried out for all four groups of mice. Consistent with previous data, CDNW/CCl_4_ mice did not develop steatosis, inflammation, and hepatocyte ballooning in 24 weeks time. Severe steatosis and steatohepatitis were observed in vehicle treated WDSW/CCl_4_ mice. Curcumin treatment drastically improved the MASH features in WDSW/CCl_4_ mouse groups and showed no and/or adverse effects on CDNW/CCl_4_ mice (**Figure 2A and 2B**). WDSW/CCl_4_ mice with vehicle control showed evident NASH development with marked steatosis (2.8±0.2), hepatocellular ballooning (1.9±0.12), and lobular inflammation (2.4±0.24) with a MASLD activity score of 7.8±0.3. As hypothesized, curcumin treatment showed a significant improvement in MASH hallmarks with a notable reduction in steatosis (1.4+/-0.2), hepatocellular ballooning (1.2±0.2), and lobular inflammation (1.4±0.24) with MASLD activity score of 3.7±0.4 in WDSW/CCl_4_ mice (**Figure 2C to 2F**). Numerous tumors (8-10 per mouse) were observed on the liver surface in 7 out of 8 mice treated with vehicle in WDSW/CCl_4_, while very few (1-2 per mouse) to no tumors in curcumin treated WDSW/CCl_4_ mouse groups (**Figure 2A, 2G and 2H**). Taken together, these data suggest that curcumin treatment improves MASH liver histology and attenuates the progression of MASH to HCC.

**Figure 2.**
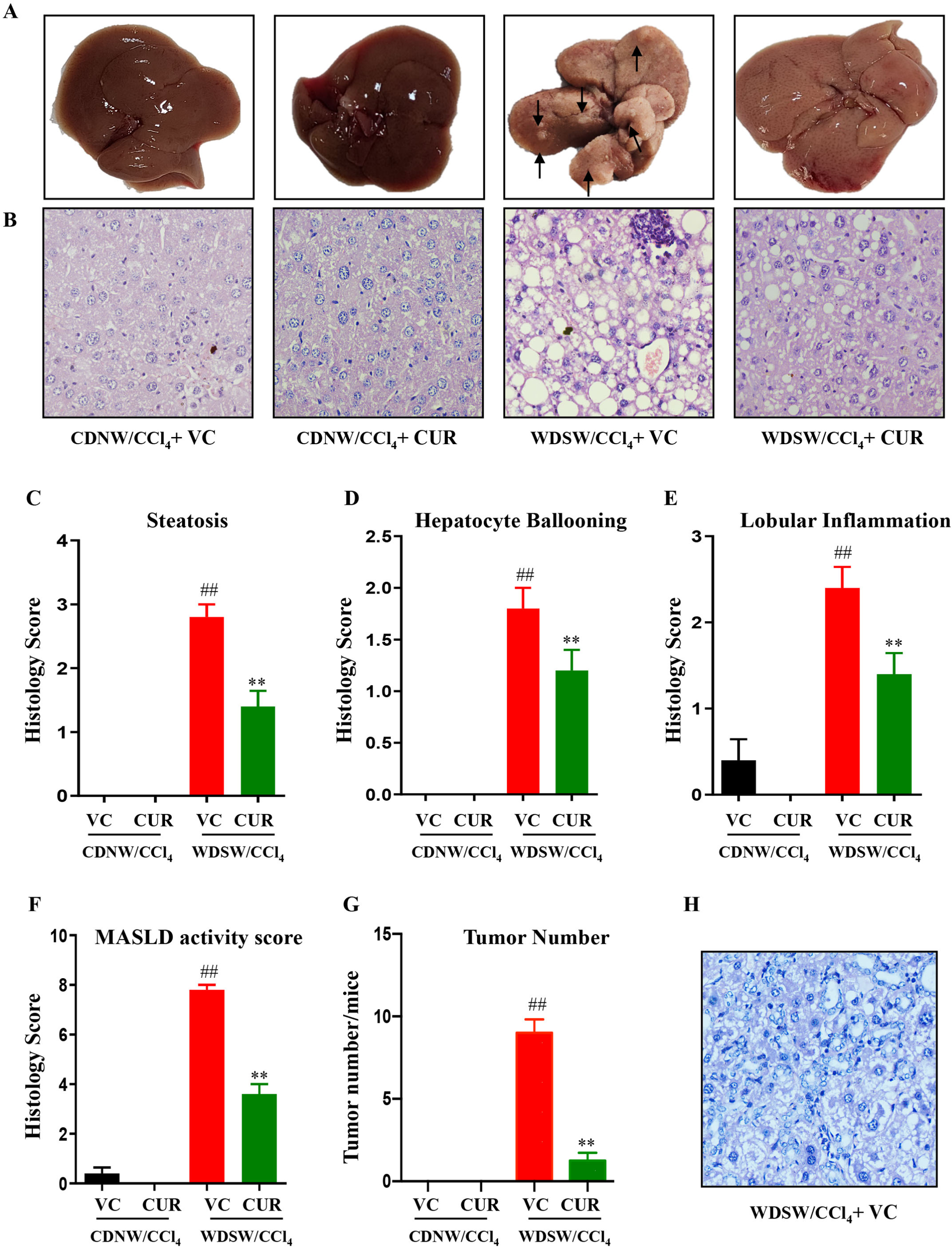
Curcumin treatment improves MASH and attenuates liver carcinogenesis. CDNW/CCl_4_ or WDSW/CCl_4_ mice were treated with vehicle (VC) or curcumin (CUR) for 12 weeks. (A) Representative images of the liver depicting the presence or absence of tumors, (B) Representative microscopic images of liver sections of each group stained with hematoxylin & eosin. Histology scores for (C) hepatic steatosis, (D) hepatocyte ballooning, (E) lobular inflammation, and (F) MASLD activity score. (G) tumor numbers and (H) tumor histology. Data are expressed as mean±SEM with 6-8 mice in each group. ## p < 0.001 or # p < 0.05 compared to CDNW/CCl_4_ with vehicle control; ** p < 0.001 or * p < 0.05 compared to WDSW/CCl_4_ with vehicle control.

### Curcumin ameliorates inflammation, fibrosis and tumorigenesis in experimental MASH-HCC

Next, we examined the effect of curcumin on liver inflammation, a key player in MASH settings. Proinflammatory cytokines such as interleukin-1β (IL-1β), interleukin-6 (IL-6), and tumor necrosis factor α (TNFα) that were overexpressed in vehicle treated WDSW/CCl_4_ mice were reduced upon the administration of curcumin, marking its anti-inflammatory effects (**Figure 3A, 3B, and 3C**). Further, the stress-responsive mitogen-activated protein kinases (MAPKs) activated due to these proinflammatory cytokines were examined. Curcumin administration downregulated the phosphorylation of extracellular signal-regulated protein kinase (ERK1/2) and c-Jun N-terminal kinase (JNK) compared to WDSW/CCl_4_ mouse groups with vehicle control (**Figure 3D and 3E**).

**Figure 3.**
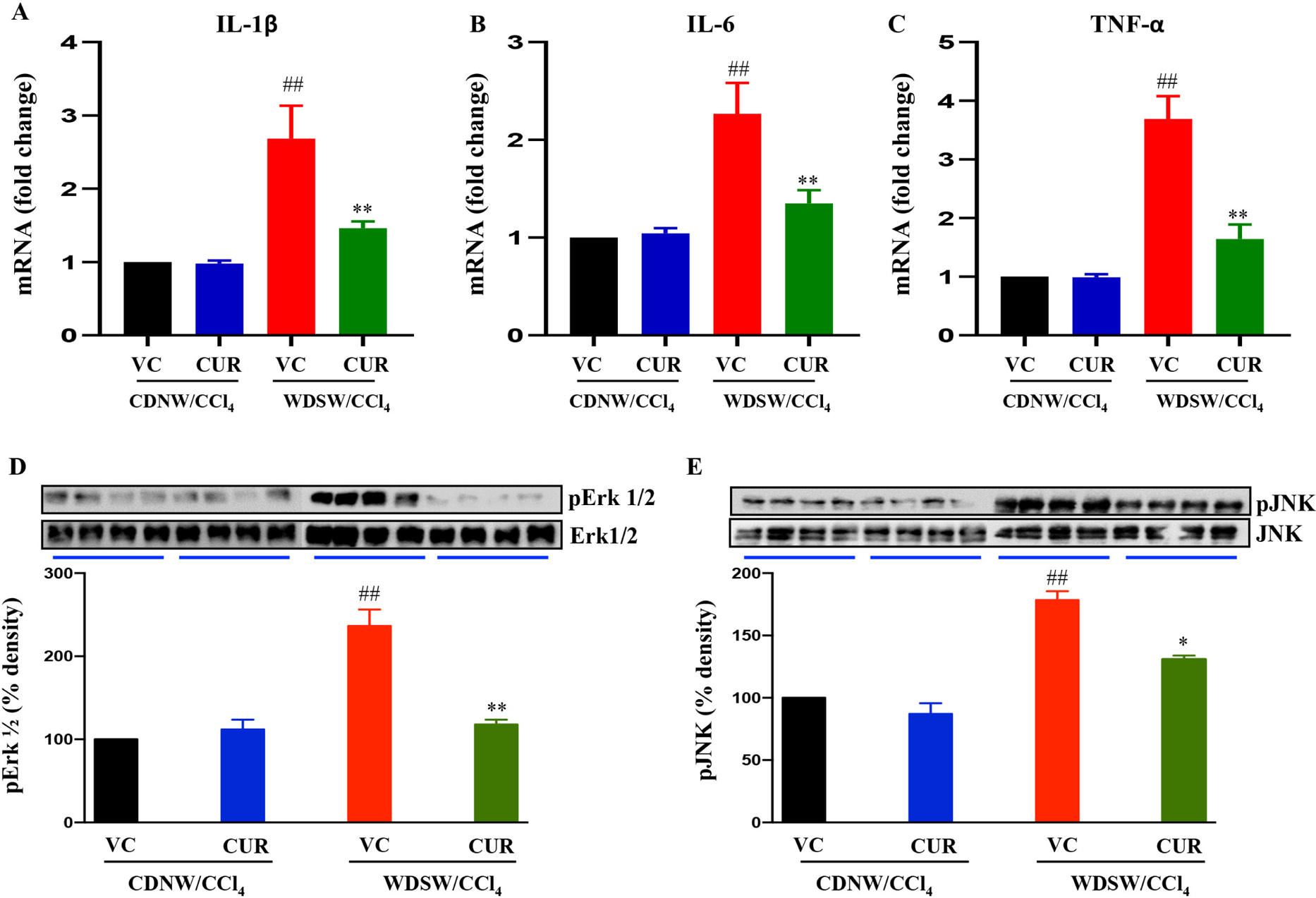
Curcumin treatment suppresses hepatic inflammation in MASH-HCC mice. The relative hepatic mRNA expression of (A) IL-1β, (B) IL-6, and (C) TNFα was determined by qRT-PCR. The expression levels were normalized with β-actin and carried out in triplicate. Immunoblots performed in the whole cell lysates of liver tissues treated with vehicle (VC) or curcumin (CUR) for (D) p-Erk1/2 and Erk1/2, (E) p-JNK and JNK. Data are expressed as mean±SEM. ## p < 0.001 or # p < 0.05 compared to CDNW/CCl_4_ with vehicle control; ** p < 0.001 or * p < 0.05 compared to WDSW/CCl_4_ with vehicle control. IL-1β, interleukin-1β; IL-6, interleukin-6; TNFα, tumor necrosis factor α; p-Erk1/2, phospho-extracellular signal-regulated protein kinase; p-JNK, phospho-c-Jun N-terminal kinase.

Sirius red staining of liver sections was performed for all the groups to examine the effect of curcumin on liver fibrosis. WDSW/CCl_4_ mice with vehicle control showed significant fibrosis, which was reduced, in curcumin-treated mice (**Figure 4A and 4B**). Further, curcumin reduced the activation of hepatic stellate cells as determined by the mRNA expression of alpha-smooth muscle actin (αSMA) (**Figure 4C**). The major fibrogenic gene, transforming growth factor β (TGF β) was also reduced in curcumin-treated WDSW/CCl_4_ mice compared to vehicle control (**Figure 4D**).

**Figure 4.**
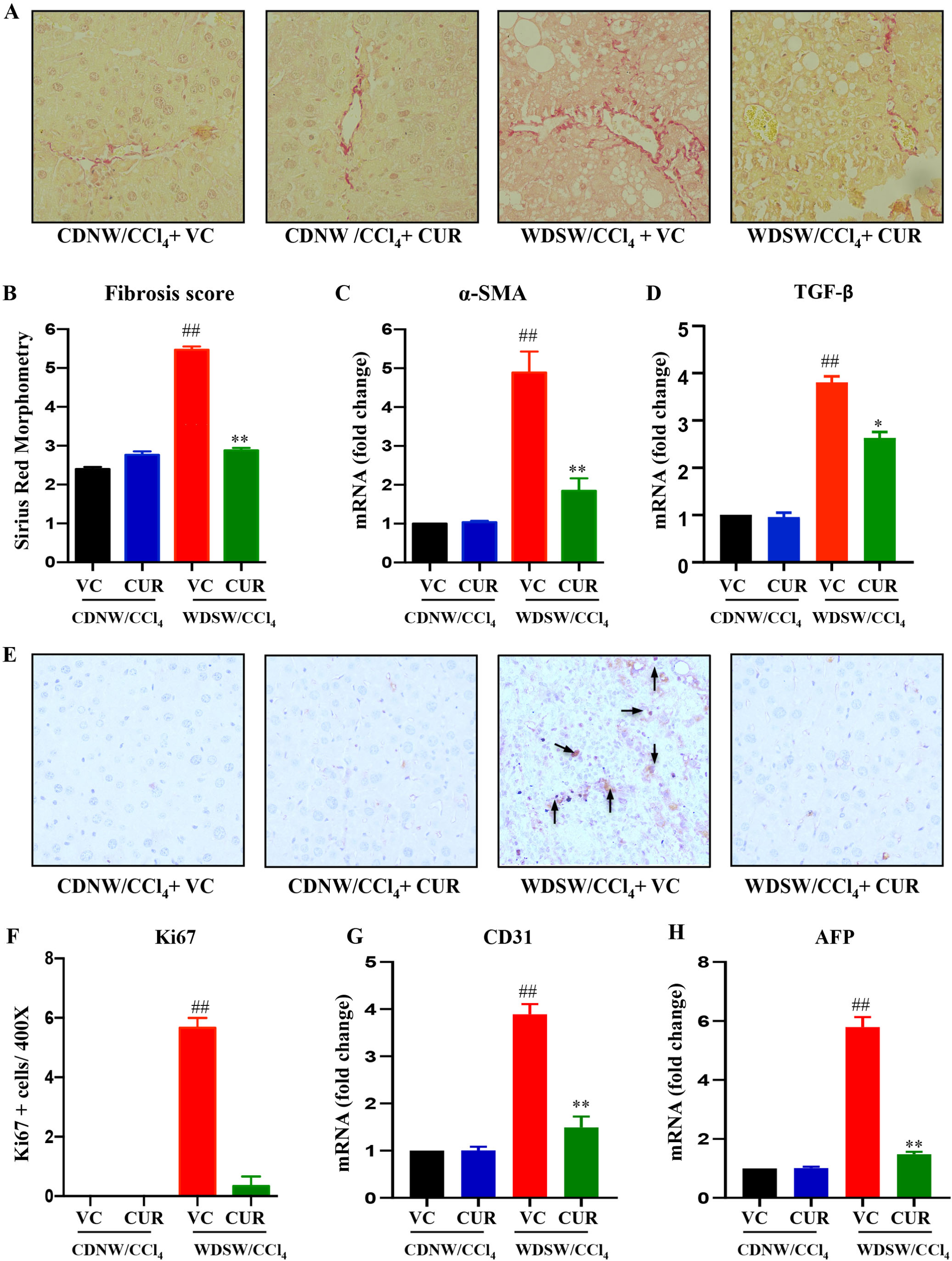
Curcumin treatment ameliorates liver fibrosis in MASH-HCC mice. CDNW/CCl_4_ and WDSW/CCl_4_ mice were treated with vehicle (VC) or curcumin (CUR) for 12 weeks. (A) Representative microscopic images of Sirius red staining of liver sections from each group. (B) Quantification of the Sirius red positive area. Hepatic mRNA expression of (C) αSMA and (D) TGF-β were estimated. Liver sections were immune stained for the (E) Ki67 marker and (F) quantified for the positive area. Hepatic mRNA expression of (G) CD31 and (H) AFP was measured by qRT-PCR. Data are expressed as mean±SEM. ## p < 0.001 or # p < 0.05 compared to CDNW/CCl_4_ with vehicle control; ** p < 0.001 or * p < 0.05 compared to WDSW/CCl_4_ with vehicle control. αSMA, alpha Smooth Muscle Actin; TGF β, transforming growth factor β; CD31, cluster of differentiation 31; AFP, alpha fetoprotein.

Additionally, we also verified the preventive effect of curcumin on liver tumorigenesis initiated by MASH by immunostaining of the proliferative marker Ki67 and observed that curcumin-treated mice livers showed negative Ki67 immunostaining against positive staining in WDSW/CCl_4_ mice with vehicle control (**Figure 4E and 4F**). We then investigated the mRNA levels of cluster of differentiation 31 (CD31), a specific marker for vascular differentiation, and alpha-fetoprotein (AFP), a key biomarker of HCC. Curcumin treatment downregulated both CD31 and AFP expression levels compared to WDSW/CCl_4_ mice with vehicle alone (**Figure 4G and 4H**). Collectively, these data provide evidence that curcumin treatment improves WDSW/CCl_4_-induced inflammation and fibrosis and prevents MASH-associated liver tumorigenesis.

### Curcumin treatment suppresses AATF-mediated MASH-HCC

Next, we wanted to understand the possible molecular targets of curcumin in preventing MASH-HCC. Owing to this objective, our previous findings have shown elevated levels of AATF in MASH-associated HCC in a diet-induced animal model of NAFLD (DIAMOND) [21]. AATF, an oncogene overexpressed in many cancers, including HCC [22], is responsible for the activation of key signaling molecules involved in tumorigenesis. Thus, in this study, we investigated the effect of curcumin on AATF levels in MASH-HCC. The immunostaining and mRNA expressions of AATF were significantly reduced upon curcumin treatment when compared to vehicle control in WDSW/CCl_4_ mouse groups (**Figure 5A and 5B**). To determine the non-toxic but effective concentrations of curcumin against human HCC cells, cell viability assay was performed. Curcumin treatment affected the cell viability of QGY-7703 cells in a dose- and time-dependent manner within 24 hours of exposure in contrast to the control group (**Figure 5C and 5D**). Furthermore, to understand the specific effect of curcumin on AATF, we used stable clones of control and AATF knockdown QGY-7703 cells. We observed that AATF expression upon curcumin treatment was reduced in control QGY-7703 cells, but no changes were seen in AATF knockdown QGY-7703 cells (**Figures 5E and 5F**). Furthermore, we evaluated the proliferative activity and migration ability of the control and AATF knockdown QGY-7703 cells upon treatment with curcumin (**Figure 5G and 5H**). Interestingly, curcumin showed a similar inhibitory effect as that of AATF knockdown cells on the proliferation and migration of human HCC cells. These results demonstrated that curcumin treatment could suppress AATF expression in MASH-HCC and thereby prevent tumorigenesis.

**Figure 5.**
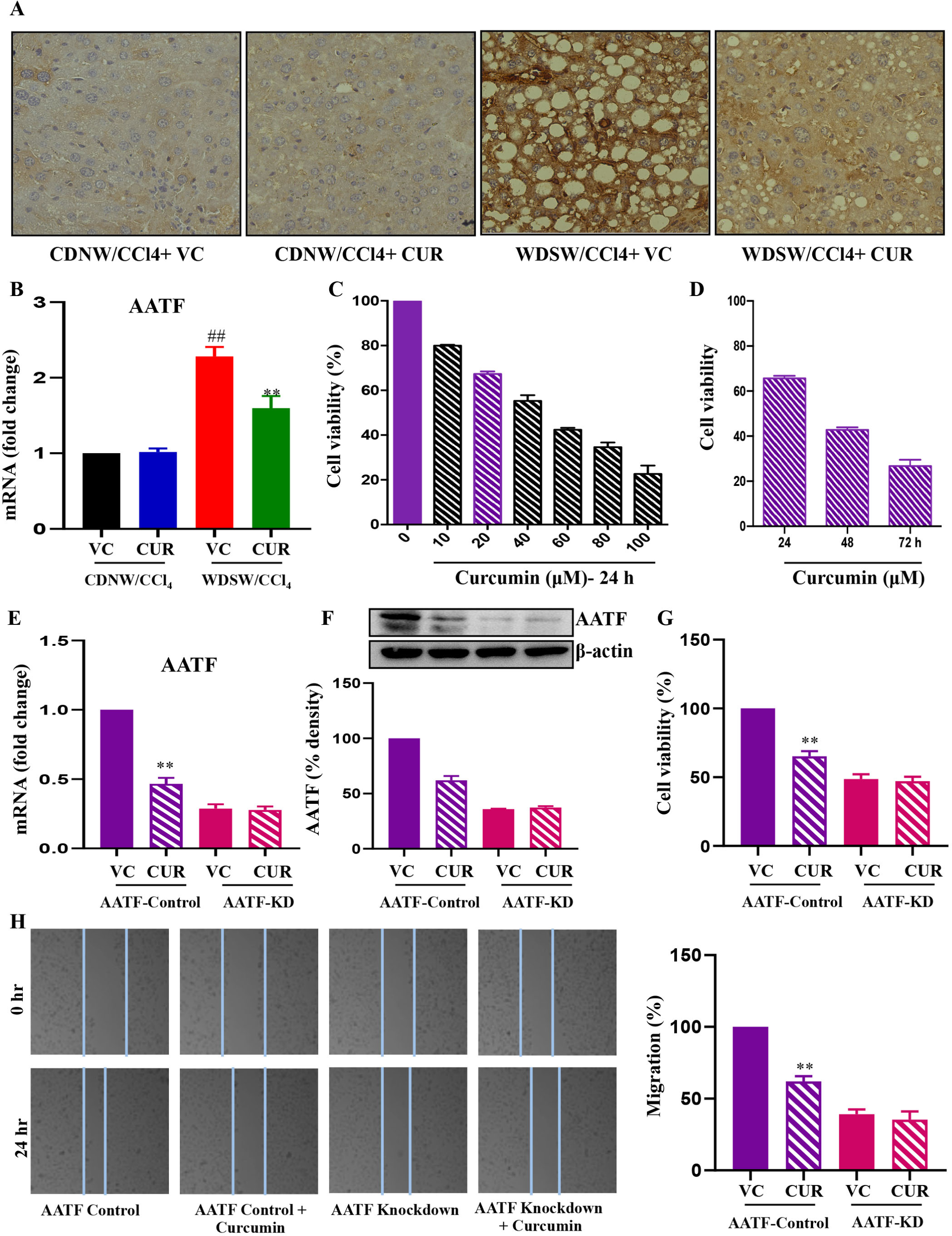
Curcumin treatment downregulates AATF expression. Liver tissues from vehicle control (VC) and curcumin (CUR) treated mice from CDNW/CCl_4_ and WDSW/CCl_4_ were processed. (A) Representative micrographs of AATF immunostaining in all four groups. (B) Relative mRNA expression of AATF in mouse liver tissues normalized to the endogenous control, β-actin. Viability of QGY-7703 cells following curcumin treatment with different concentrations (C) and time (D). The mRNA expression (E) and immunoblot (F) of AATF in control and AATF knockdown (KD) QGY-7703 cells treated with vehicle (VC) or curcumin (CUR). (G) Cell viability of control and AATF knockdown (KD) QGY-7703 cells treated with vehicle (VC or curcumin (CUR) normalized against control. (H) Representative micrographs (100X) and quantification of cell migration of control and AATF knockdown (KD) QGY-7703 cells treated with vehicle (VC) or curcumin (CU). Data are expressed as mean±SEM. ## p < 0.001 or # p < 0.05 compared to CDNW/CCl_4_ with vehicle control; ** p < 0.001 or * p < 0.05 compared to WDSW/CCl_4_ with vehicle control or AATF control cells treated with vehicle.

### Molecular mechanism of AATF inhibition by Curcumin

Further, we sought to explore the possible mechanisms of curcumin-mediated downregulation of AATF. Previous studies have demonstrated that curcumin treatment demethylates the CpG islands of the promoter of KLF4, increasing its expression [36]. Thus, we evaluated the expression of KLF4 in the mouse liver tissue of all four groups. Interestingly, KLF4 was upregulated in curcumin-treated WDSW/CCl_4_ mice in contrast to vehicle alone (**Figure 6A and 6C**). KLF4, being a tumor suppressor gene, is implicated in the downregulation of several oncogenes in HCC. Studies have shown that KLF4 negatively regulates the expression of Sp1, a pro-oncogenic molecule that binds directly to the AATF promoter and induces its expression [37,16]. Following this hypothesis, in our study, curcumin treatment showed a marked reduction in Sp1 levels in the WDSW/CCl_4_ mouse groups compared to vehicle control (**Figure 6B and 6C**). Similarly, the human HCC cells, QGY-7703 cells upon treatment with curcumin also showed upregulated levels of KLF4 (**Figure 6D and 6E**). This was supported by the downregulation of Sp1 in the QGY-7703 cells treated with curcumin (**Figures 6F and 6G**). Overall, these data indicate that curcumin via KLF4 inhibits the expression of Sp1, which in turn is responsible for the downregulation of AATF. Next, we evaluated the levels of activated STAT3, a prominent transcription factor that plays a key role in the progression and metastasis of HCC [38]. As anticipated, the pSTAT3 levels were downregulated in curcumin treated WDSW/CCl_4_ mice compared to the vehicle control (**Figure 6H).** Similarly, pSTAT3 levels were reduced in curcumin-treated control QGY-7703 cells in contrast to AATF knockdown cells (**Figure 6I**). Overall, the data indicate that curcumin-mediated AATF inhibition prevents the disease progression of MASH to HCC.

**Figure 6.**
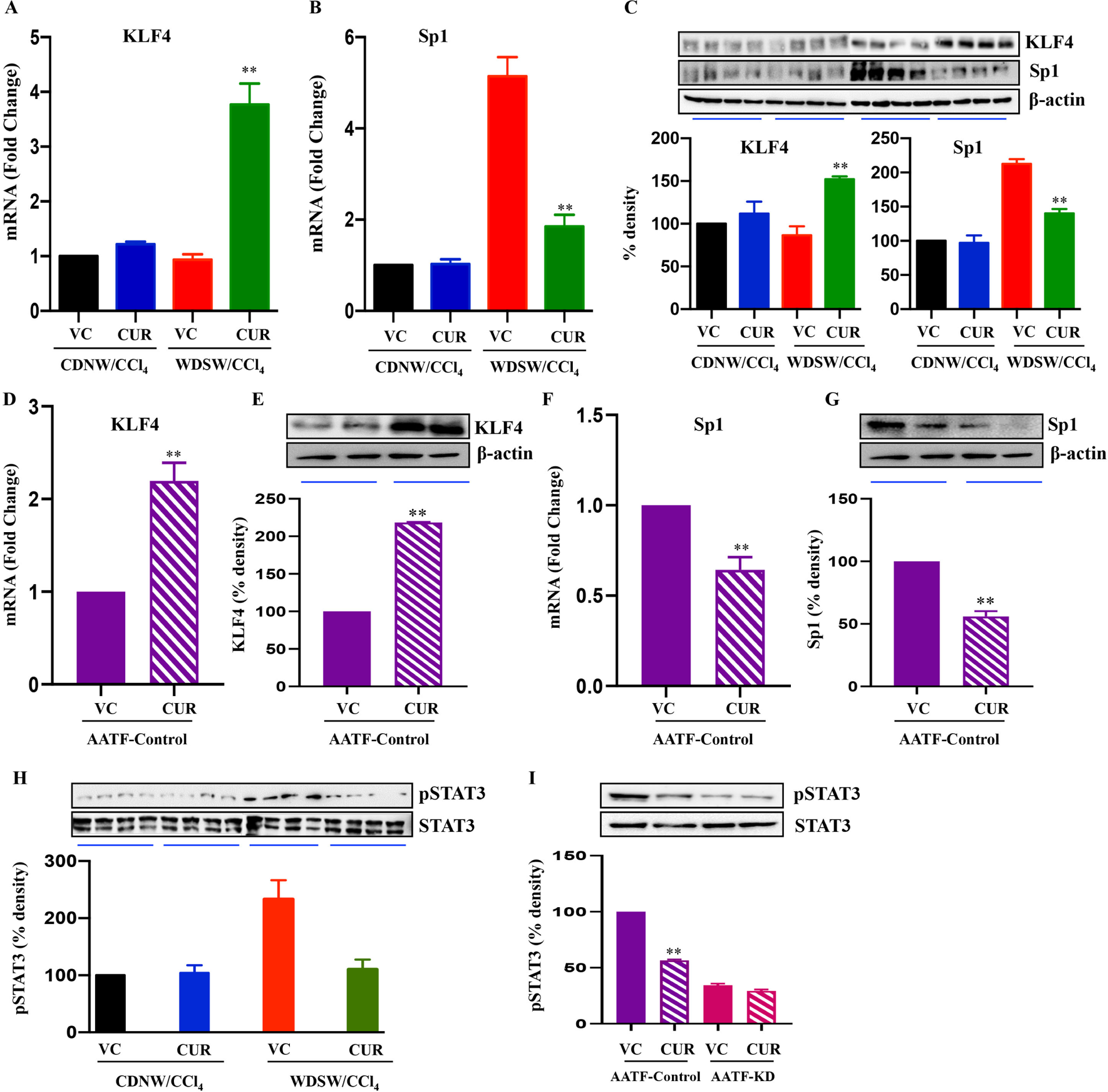
Curcumin downregulates AATF via KLF4 and Sp1 signaling pathway. The relative mRNA expression of KLF4 (A) and Sp1 (B) and (C) immunoblots of KLF4 and Sp1 normalized to the endogenous control β-actin from CDNW/CCl_4_ and WDSW/CCl_4_ mice treated with vehicle (VC) or curcumin (CUR). QGY-7703 cells treated with vehicle (VC) or curcumin (CUR) were examined for the mRNA and protein expression of KLF4 (D and E) and Sp1 (F and G). Whole cell lysates prepared from the control and AATF knockdown (KD) QGY-7703 cells treated with vehicle (VC) or curcumin (CUR) were also checked for the expression of pSTAT3 and STAT3, pSTAT3 normalized to its control STAT3. Data are expressed as mean±SEM. ** p < 0.001 or * p < 0.05 compared to WDSW/CCl_4_ with vehicle control or AATF control cells treated with vehicle. KLF4, kruppel-like factor 4; Sp1, specificity protein 1.

**Figure 7.**
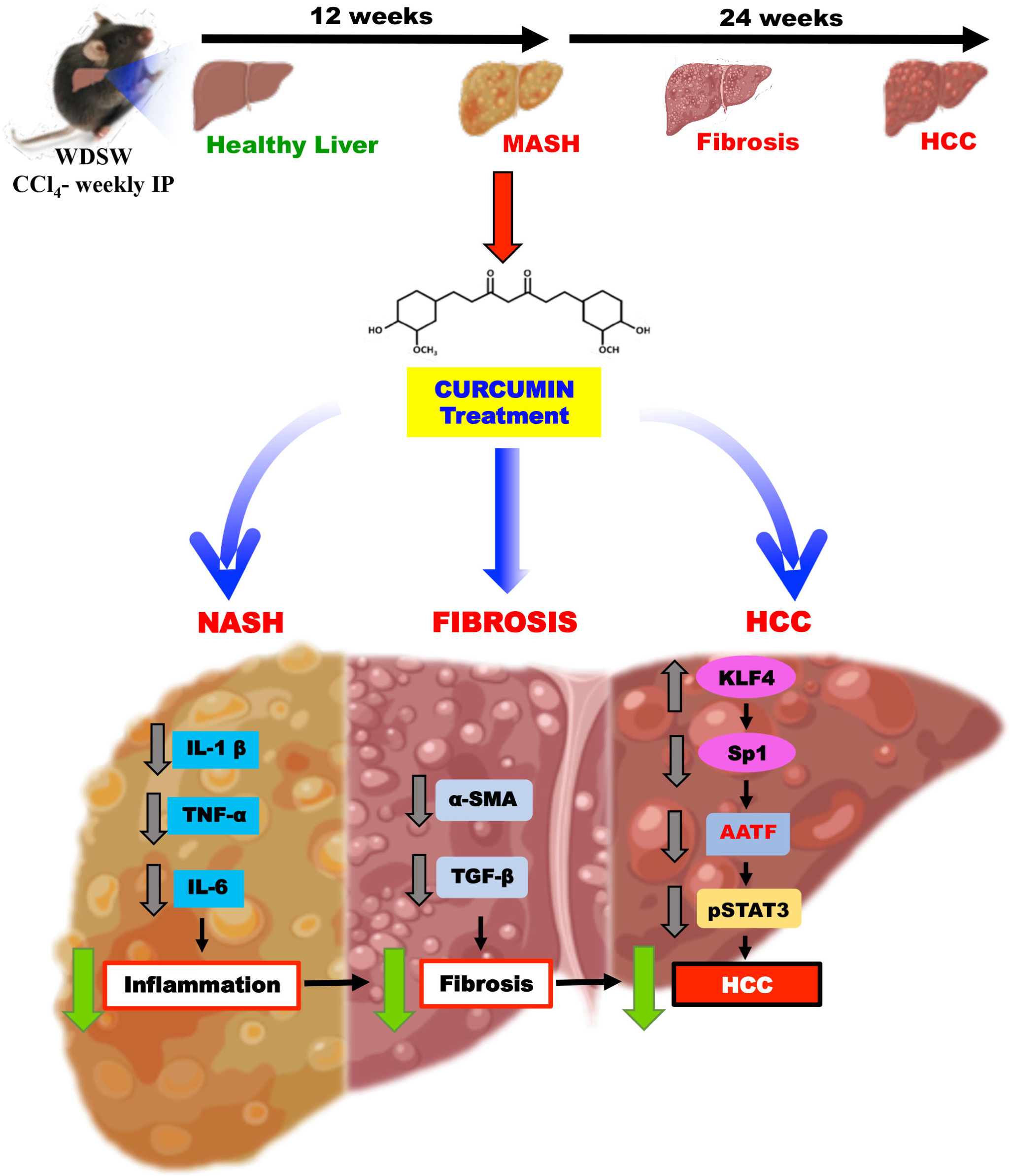
Molecular mechanisms involved in the therapeutic action of curcumin in ameliorating AATF-mediated MASH-HCC. CDNW/CCl_4_ and WDSW/CCl_4_ were treated with curcumin for 12 weeks. Administration of curcumin ameliorated steatohepatitis (MASH), fibrosis and progression to HCC in WDSW/CCl_4_ mice by downregulating AATF expression via KLF4 and Sp1 signaling pathway.

## Discussion

HCC, a type of liver malignancy, has a high mortality rate, and current treatment options are limited and often ineffective [39]. MASLD is the fastest-growing cause of the rising prevalence of HCC. Apart from the linear progression of MASH to fibrosis, cirrhosis, and HCC, recent studies have shown the occurrence of HCC in patients without cirrhosis as well [3]. The complex process of MASH progressing in HCC is linked to the metabolic, immunological, and inflammatory milieu induced by insulin resistance and lipotoxicity [8]. Due to poor prognosis and limited studies on molecular pathogenesis, MASH-associated HCC management is a challenge. There are no approved drugs to treat MASH-HCC despite it being a leading etiology for oncogenesis. Added to this, MASH-HCC is a difficult disease to treat, and patients often have poor outcomes due to late diagnosis and limited treatment options [11]. Therefore, there is a critical need for the development of effective therapeutic agents to target MASH-HCC. These agents should be able to specifically target cancer cells and minimize damage to healthy cells, thereby potentially improving outcomes, increasing survival rates, and reducing the burden of this devastating disease.

There are a number of natural therapeutic agents that have been explored for the treatment of HCC [40]. Natural therapeutic agents, such as herbs, extracts, or dietary supplements, have gained significant attention in the treatment of HCC due to their potential anti-cancer properties [41,42]. Using natural therapeutic agents for treatment can be beneficial for several reasons. Firstly, natural agents are typically less toxic than conventional chemotherapeutic drugs, which can cause significant side effects. Secondly, natural agents can be used in combination with traditional treatments such as surgery, radiation, and chemotherapy to enhance their efficacy and reduce side effects. Finally, natural agents are often more affordable and accessible than conventional drugs, making them an attractive option for patients who cannot afford expensive treatments. Overall, the use of natural therapeutic agents in the treatment of HCC can be a safe and effective option for patients looking to manage their cancer symptoms and improve their overall quality of life. Along the same lines, curcumin is one such naturally occurring compound that is well known for its efficacy and safety [26]. It has been studied for its potential health benefits, and research suggests that it may have a variety of targets in the human body. Some of the targets of curcumin include inflammation, oxidative stress, cancer, brain function, and heart health [27,43,44]. There is growing evidence from both preclinical and clinical studies that suggests curcumin may have anti-cancer effects against HCC by exerting anti-inflammatory effects, apoptosis, and anti-angiogenic effects [30,45,46]. However, less is explored in the area of MASH-HCC. Studies have suggested that curcumin may help reduce liver inflammation and oxidative stress, which are key factors in the development and progression of MASH [10]. Additionally, curcumin has been found to have beneficial effects on lipid metabolism, insulin resistance, gut microbiota, and fibrosis, all of which are associated with MASH and potentially the progression of MASH to HCC [5]. Nonetheless, additional validation is required to confirm the potential therapeutic efficacy of curcumin in inhibiting AATF-mediated MASH-HCC.

There are several murine models of MASH and MASH-HCC, but most do not recapitulate the human disease, which is a major drawback [47]. In this study, we have used a murine model of MASH that could be induced with a simple diet and chemical, which progresses through the stages of steatosis, steatohepatitis, fibrosis, and liver cancer [32]. Of note, this experimental model employs intraperitoneal injection of a very low dose of CCl_4_ (0.2 μl/g) once per week, starting simultaneously with diet administration. The WDSW consists of a high fat, high sucrose, and high cholesterol diet (WD, 21% fat, 41% sucrose, and 1.25% cholesterol by weight) with sugar water (SW, 18.9 g/L d-glucose and 23.1 g/L d-fructose). The disease development in the WDSW/CCl_4_ FAT-NASH (**F**ibrosis **A**nd **T**umors) murine model closely resembles the metabolic, histological, and transcriptomic signature of human MASH-HCC, making the study more significant and translational. Herein, we evaluated the hepatoprotective effect of curcumin using this experimental MASH-HCC model. Interestingly, we found that mice treated with vehicle in WDSW/CCl_4_ for 24 weeks readily developed HCC with numerous visible tumors on the liver. Previously, studies by Afrin R et al., demonstrated that curcumin would reduce the progression of MASH to HCC by inhibiting the cytosolic translocation of high mobility group box 1 (HMGB1) and the nuclear translocation of NF-κB using a streptozotocin-HFD-induced MASH-HCC mouse model [30]. However, artificial induction of MASH by streptozotocin and lack of translational relevance in various aspects restrict the ability to fully evaluate the therapeutic effects of curcumin in this particular model. Curcumin treatment in our study attenuated tumor formation by improving WDSW-induced obesity and liver damage. The treatment of curcumin to the WDSW/CCl_4_ mice resulted in an improvement in obesity, as marked by a reduction in body weight and liver weight. Furthermore, the study demonstrated amelioration in insulin resistance and marked improvement in the biochemical and histological outcomes after treatment with curcumin in the WDSW/CCl_4_ mouse groups. Studies have reported that inflammatory cytokines TNF-α, and IL-6 targeting JNK, ERK, and NF-κB pathways drive HCC arising in inflammatory settings [10,48]. Curcumin treatment successfully lessened the inflammatory cytokines as well as stress kinases. It is well established that inflammatory setting gives rise to or predisposes to fibrosis and HCC. The cytokine-driven interactions between the stellate cells and the creation of the microenvironment lead to the emergence of HCC development [49]. In this regard, in our experiments, we found that the elevated expression of fibrogenic genes and oncogenic markers in WDSW/CCl_4_ mice was reduced upon treatment with curcumin. Collectively, our study reports the effect of curcumin on attenuating the progression of MASH to HCC while also improving the existing MASH condition.

The current study provides insight into the mechanism by which curcumin prevents MASH-associated hepatocarcinogenesis mediated by AATF. Curcumin has been found to inhibit the activation of oncogenes and also induce apoptosis in HCC cells. It has also been shown to have anti-inflammatory effects, which can help prevent the development of HCC [24,43]. Furthermore, we examined the effect of curcumin on AATF, an oncogene that is overexpressed in MASH-HCC. Interestingly, our data showed that curcumin treatment downregulated AATF expression in mouse liver tissues. In order to elucidate the molecular mechanisms of how curcumin acts on AATF, stable clones of AATF control and AATF knockdown were prepared in human HCC cells, QGY-7703. The oncogenic properties of the QGY-7703 cells, such as increased cell proliferation, and cell migration were inhibited upon curcumin treatment in control cells and had no effect on the AATF knockdown cells. Studies by Chatterjee B. et al., have demonstrated that KLF4, a key player in regulating cell growth, differentiation, and development, is activated by curcumin. Demethylation of CpG islands on the KLF4 promoter by curcumin is known to increase its expression and also its promoter occupancy to p21, leading to curcumin-mediated cell cycle arrest and inhibition of cell proliferation [36]. Similarly, our studies showed a significant increase in the expression of KLF4 in WDSW/CCl_4_ mice treated with curcumin compared to vehicle controls. Consistently, upon curcumin treatment, there was upregulation of KLF4 in AATF control QGY-7703 cells. In another study by Kanai M. et al., loss of KLF4 leads to sp1 overexpression and promotes the development and progression of human gastric cancer [37]. Thus, KLF4 negatively regulates the expression of Sp1, a pro-oncogenic molecule that binds directly to the AATF promoter and induces its expression. Consistent with these studies, we found that in WDSW/CCl_4_ mice and AATF control cells, curcumin treatment reduced AATF expression by downregulating Sp1 via KLF4 and thereby exerting anti-tumor activity. It is reported that STAT3 plays a key oncogenic role in cell proliferation, migration, immune suppression, and the angiogenesis of HCC [38]. Studies by Kumar et al., have shown that AATF drives HCC via monocyte chemoattractant protein 1 (MCP1) and STAT3 [21]. Consistently, p-STAT3 levels were reduced by curcumin treatment in WD/CCl_4_ mice and AATF control HCC cells but had no effect on AATF knockdown cells.

In conclusion, the current study adds to the growing body of evidence supporting the promising role of curcumin and its potential applications in the prevention and treatment of MASH-HCC. We demonstrated that curcumin ameliorates AATF-mediated liver damage and inflammation to cancer in MASH-HCC via KLF4 and Sp1 signaling pathway. Furthermore, the study offers valuable insight into the potential benefits of curcumin for MASH-HCC, for which the development of effective therapeutic agents is an absolute necessity, and additionally provides a basis for more rigorous clinical trials to fully evaluate the effectiveness of curcumin for MASH-HCC.

## Abbreviations

HCC: hepatocellular carcinoma
MASLD: metabolic dysfunction associated steatotic liver disease
NAFLD: nonalcoholic fatty liver disease
MASH: metabolic dysfunction associated steatohepatitis
NASH: nonalcoholic steatohepatitis
AATF: apoptosis antagonizing transcription factor
STAT3: signal transducer and activator of transcription 3
TNF-α: tumor necrosis factor-α
ER: endoplasmic reticulum
NF-κB: nuclear factor kappa B
KLF4: kruppel-like factor 4
Sp1: specificity protein 1
CDNW: chow diet normal water
WDSW: western diet sugar water
CCl_4_: carbon tetrachloride
VC: vehicle control
ALT: alanine transaminase
AST: aspartate transaminase
IL-1β: interleukin-1β
IL-6: interleukin-6
ERK1/2: extracellular signal-regulated protein kinase
JNK: Jun N-terminal kinase
CD31: cluster of differentiation 31
AFP: alpha-fetoprotein

## Author Contributions

Akshatha N. Srinivas and Diwakar S.: Writing-original draft, Investigation, Methodology, Formal Analysis, Visualization. Saravana Babu Chidambaram: Writing-review & editing. Prasanna K. Santhekadur: Formal analysis, Writing-review & editing. Divya P. Kumar-Funding acquisition, Supervision, Conceptualization, Project administration, Formal analysis, Writing-original draft.

## Acknowledgements

We acknowledge the infrastructure support grant provided by the Department of Science & Technology to the CEMR Laboratory (CR-FST-LS-1/2018/178); to Department of Biochemistry (SR/FST/LS-1-539/2012) at JSS Medical College, JSS Academy of Higher & Research, Mysuru, Karnataka, INDIA.

